# Reducing Sanger Confirmation Testing through False Positive Prediction Algorithms

**DOI:** 10.1101/2020.04.30.066159

**Authors:** James M. Holt, Melissa Wilk, Brett Sundlof, Ghunwa Nakouzi, David Bick, Elaine Lyon

**Author notes:** Correspondence to: James M. Holt, HudsonAlpha Institute for Biotechnology, 601 Genome Way, Huntsville, AL, USA.

## Abstract

**Purpose:** Clinical genome sequencing (cGS) followed by orthogonal confirmatory testing is standard practice. While orthogonal testing significantly improves specificity it also results in increased turn-around-time and cost of testing. The purpose of this study is to evaluate machine learning models trained to identify false positive variants in cGS data to reduce the need for orthogonal testing.

**Methods:** We sequenced five reference human genome samples characterized by the Genome in a Bottle Consortium (GIAB) and compared the results to an established set of variants for each genome referred to as a ‘truth-set’. We then trained machine learning models to identify variants that were labeled as false positives.

**Results:** After training, the models identified 99.5% of the false positive heterozygous single nucleotide variants (SNVs) and heterozygous insertions/deletions variants (indels) while reducing confirmatory testing of true positive SNVs to 1.67% and indels to 20.29%. Employing the algorithm in clinical practice reduced orthogonal testing using dideoxynucleotide (Sanger) sequencing by 78.22%.

**Conclusion:** Our results indicate that a low false positive call rate can be maintained while significantly reducing the need for confirmatory testing. The framework that generated our models and results is publicly available at https://github.com/HudsonAlpha/STEVE.

## INTRODUCTION

Clinical next-generation sequencing (NGS) is widely used to identify a molecular diagnosis in patients with suspected genetic disorders.^1,2^ Because the reported variants can impact patient care, the American College of Medical Genetics (ACMG) and the College of American Pathologists (CAP) recommends orthogonal confirmation (e.g. Sanger sequencing) for reported variants to reduce the risk of false positive results.^3,4^ Unfortunately, orthogonal confirmation increases both the cost and turn-around time of the NGS test. Furthermore, the total number of variants that are candidates for clinical reporting is steadily increasing, as demonstrated by the growth in public databases such as ClinVar and OMIM.^5,6^ Orthogonal confirmation of all reported variants will cause the effective cost of NGS to steadily increase due to an increase in the number of variants sent for confirmation.

To address this issue, other studies have questioned the necessity of orthogonal testing, especially when the variant call is of sufficiently high quality for the particular NGS assay.^7–10^ Most of these studies involved a relatively small sample size (<8000 variants), with the notable exception of the work by Lincoln et al. which examined approximately 200,000 variants identifying 1,662 as false positives.^10^ Lincoln et al. used a combination of reference samples characterized by the Genome in a Bottle Consortium (GIAB)^11–13^ along with orthogonal test results from over 80,000 clinical tests from two different laboratories. Briefly, their method involved manual selection of thresholds for quantitative metrics. These metrics were converted into a set of flags for a heuristic algorithm to classify variant calls as true positive calls or false positive calls. The exact set of flags was notably different for single nucleotide variants (SNVs) and insertions/deletions (indels). The authors establish a 100% capture rate (confidence interval 98.5%-100% for SNVs, 99.1%-100% for indels) for false positive calls while maintaining relatively low rates of calls that were incorrectly labeled as false positive calls (13.2% for SNVs, 15.4% for indels).^10^

Despite the success of the Lincoln et al. approach, there are some drawbacks to a broad application of their method. First, there are relatively few false positive variant calls (1,662 out of 200,000 variants) across their dataset, a point that is reflected in the confidence intervals for the capture rate. Second, the selection of flags from quality metrics is a manual step. This requires expert knowledge about each quality metric for a particular assay and/or pipeline, and it reduces the metric from a numerical value to a single flag value (e.g., a boolean value) which can result in a loss of information. Finally, while the majority of their orthogonal results were from GIAB, the study, nevertheless relies on a relatively large number of orthogonally confirmed results performed by the laboratory as part of clinical NGS testing (>80,000). For many labs, this is impractical due to costs, especially when developing a new test where orthogonal results are not already known.

To address difficulties associated with Lincoln’s approach we applied an automated machine learning approach^14,15^ that uses the entirety of the GIAB truth sets as the training and testing sets. The benefits of this approach are three-fold: 1) automation of quality metric evaluation on non-boolean values (i.e., no manually identified flags); 2) a substantial increase in the number of true positive and false positive variant calls available for training and testing (~3.2-3.5 million true positives per sample); and 3) no orthogonal testing results required for training. This framework, Systematic Training and Evaluation of Variant Evidence (STEVE), allows for the development of lab-specific models applicable to specific tests while permitting customization of the sensitivity settings to suit the requirements of the test.

## MATERIALS AND METHODS

### Overview

We performed clinical genome sequencing on the following Genome in a Bottle Consortium samples with published truth sets: HG001-HG005.^11–13^ These sequence data were processed using two different secondary pipelines: Illumina’s Dragen Germline Pipeline^16^ and a pipeline consisting of alignment with Sentieon^17^ and variant calling with Strelka2.^18^ Each pipeline performed both alignment and variant calling to produce a Variant Call Format (VCF) file. Each VCF file was compared to the corresponding truth set to classify each variant call as a true positive call or a false positive call. Quality metrics for each variant call were extracted directly from the VCF file and converted into machine learning features. The variant calls were divided into six distinct datasets based on the variant type and genotype of the call. The six datasets were (1) SNV heterozygotes (2) SNV homozygotes, (3) SNV complex heterozygous (two different non-reference alleles), (4) indel heterozygotes, (5) indel homozygotes and (6) indel complex heterozygous (two different non-reference alleles). Each dataset was used separately for training and testing of a machine learning model for that particular data type leading to six distinct models per pipeline, for a total of 12 models with the two pipelines we evaluated.

For each dataset, we generally followed standard machine learning practices to create our models. We tested multiple freely available algorithms to train our models. The process included splitting the dataset into training and testing sets, cross-validation, hyperparameter tuning, and a final evaluation on the testing set.^14,15^ We developed a set of clinical criteria required to pass a model, and developed a tie-breaking scheme when multiple models for a single dataset were acceptable. Subsequently, we performed a retrospective analysis on a collection of variants that had been orthogonally confirmed. Finally, we report on the clinical application of these models for non-actionable variants.

### Dataset Generation

All training and testing datasets for the machine learning models are derived from the five, well-studied Genome in a Bottle (GIAB) samples.^11–13^ Briefly, these samples consist of NA12878 (HG001), a well-studied female of European ancestry; HG002-004, a trio (son and parents) of Ashkenazi Jewish ancestry; and HG005, a male of Chinese ancestry. GIAB provides high-confidence call regions, variants within those regions, and genotype calls for each variant for each of these five samples. Each high-confidence region covers 80-90% of reference genome hg38 and contains approximately 3.2-3.5 million, non-reference variant genotypes for the corresponding sample.

DNA was purchased from Coriell or NIST (see Supplemental Materials) and sequenced with the NovaSeq 6000 sequencing platform. The DNA was sonicated and prepared as a paired-end library with ligation of Illumina flowcell-specific adapter sequences and a unique barcode. The prepared library was then quality checked for adequate yield through fluorescence methods and quantitative polymerase chain reaction (PCR), as well as for appropriate library size and profile using bioanalysis. Libraries were clustered onto Illumina NovaSeq 6000 flowcells and sequenced using standard Illumina reagents and protocols. The output of this protocol is paired-end 150bp reads in FASTQ format with a mean coverage of at least 30x and passing stringent quality control metrics.

The data was aligned to the human reference genome (hg38) and variants were called using Illumina’s Dragen Germline Pipeline.^16^ Alignment and variant calling was also performed using a second pipeline consisting of alignment with Sentieon and variant calling with Strelka2.^17,18^ The output of each pipeline consisted of a single Variant Call Format (VCF) file. These VCF files were matched with the corresponding GIAB high-confidence regions and call sets and evaluated using the Real Time Genomic (RTG) VCFeval tool.^19^ VCFeval is capable of handling differences in variant representation and genotype differences while restricting the evaluation to only the high-confidence regions. The final output consists of two VCF files per sample-pipeline combination: one containing all variants labeled as true positive calls and one containing all variants labeled as false positive calls.

We then converted the VCF files into machine learning labels and features. First, labels were assigned based on the RTG VCFeval output file. All variants in the true positive file were labeled as true positives, and all variants in the false positive file were labeled as false positives. Features were extracted directly from the VCF files as well. Generally, these were numerical values corresponding to quality metrics generated by the upstream pipeline. Importantly, the set of quality metrics available from each pipeline were different and shared metrics may be calculated differently due to implementation differences. Thus, the data from each pipeline was handled independently to create pipeline-specific models. We detail the precise set of features extracted for each pipeline in the Supplemental Material.

For each pipeline dataset, we stratified all of the labels and features into one of six machine learning datasets based on the variant type and genotype combination. We used two categories for variant type (SNV or indel) and three categories for genotype (heterozygous, homozygous, and complex heterozygous (two different non-reference alleles)). Each of the six datasets was handled independently using an identical process that is detailed in the following section.

### Model Training and Testing

Our primary goal was the accurate identification of false positive variant calls. We also sought to minimize the number of true positive calls that would be labeled *incorrectly* as false positive calls. Since false positive variant calls (variants called by the pipeline but absent from the truth set) are the primary target, they are labeled as positives (binary label “1”) when passed to the machine learning algorithms. Similarly, true positive variant calls passed to the machine learning algorithm are labeled as negatives (binary label “0”). The goal of machine learning in this application is to create a model with high sensitivity, meaning that few or no false positive variant calls will be missed by the model and allowed onto the final patient report. The model should also have a high specificity, meaning that the lowest number of true positive variant calls will be flagged by the model to be sent for confirmatory testing.

In general, we followed the machine learning guidelines recommended by Scikit-Learn (sklearn).^20^ We first split the variants from each sample into equal sized training and testing datasets such that the number of false positive and true positive variant calls were balanced. The testing dataset was set aside and only used in the final evaluation.

We selected four algorithms for model generation that each conforms to the sklearn paradigm: AdaBoost, EasyEnsemble, GradientBoosting, and RandomForest.^21–24^ We also selected hyperparameters for each model which were automatically evaluated during cross-validation (see below). See the Supplemental Material for further details concerning the hyperparameters evaluated.

We performed a leave-one-sample-out cross validation using the training data.^14^ Given *S* samples, the models are trained on (*S*-1) samples then evaluated using the left-out sample to simulate receiving a “new” sample. This is performed a total of *S* times (each sample is left out once), leading to a 7-fold, leave-one-sample-out cross validation in our analysis. As noted earlier, hyperparameters were automatically tested during the cross-validation process and the best performing hyperparameters (based on Area Under the Receiver-Operator Curve) were used during the final training process.

Additionally, each model was evaluated at eight different sensitivity values in the range of 99%-100%. These “evaluation” sensitivities represent different thresholds that a clinical laboratory might select as a requirement for their test. 99% represents a sensitivity that is likely at the lower end of acceptable practice (i.e. 1/100 false positive calls are missed) whereas 100% sensitivity (i.e. *no* false positive calls missed) represents a clinical goal that is desirable but rarely achievable in practice. With six variant/call combinations, seven leave-one-sample-out cross-validation evaluations, and 45 model/hyperparameter combinations, a total of 1890 models were trained during this process.

The final step of the process is to re-train the models using only the best hyperparameters for each model and the full training set. Once trained, the models were then evaluated on the testing dataset that was previously set aside. As noted earlier, each of these models was evaluated at eight different evaluation sensitivities, leading to a total of 32 candidate hypertuned models for each variant/genotype dataset.

### Clinical Application

After the algorithms were trained, we developed a set of criteria to identify an algorithm to introduce into clinical practice. Given the results from the 7-fold cross-validation, we calculated both mean and standard deviation of each model’s sensitivity. We defined the lower bound of sensitivity as two standard deviations below the mean (−2SD) and the upper bound as two standard deviations above the mean (+2SD).

We then selected both a minimum acceptable sensitivity and a target sensitivity (i.e. the desired sensitivity). Given those two values, we enforced two criteria for a model to pass: 1) the lower bound of the cross-validation sensitivity (−2SD) must be greater than or equal to the minimum acceptable sensitivity and 2) the final testing sensitivity must be within the bounds of the cross-validation sensitivity ([−2SD, +2SD]). The first requirement provides confidence that the trained models are consistently performing above the minimum acceptable sensitivity. The second requirement provides confidence that the final trained model is consistent with the results from cross-validation and helps reject final models that are suffering from overfitting or underfitting. Because we had multiple evaluation sensitivities, several models passed these two criteria. In order to break ties, we developed a modified F1 score that incorporates both the sensitivity and specificity of the models in order to choose a single model for clinical practice. Details of this implementation along with results from the trained models can be found in the Supplemental Material.

Given a set of accepted clinical models (one per variant/genotype combination), we used the models to perform a retrospective analysis of orthogonally confirmed variants that had been previously reported by the HudsonAlpha Clinical Services Lab (CSL). All variants were chosen from cGS cases reported by the CSL between October 2, 2019 and December, 11 2019. The variants chosen contained a mixture of primary findings, actionable secondary findings, carrier status findings, and pharmacogenomic findings. Each variant is associated with a VCF file that was generated using an identical process as the VCFs used in the model training. Finally, we report the results of this approach in clinical practice, applied to carrier status findings and pharmacogenomic findings. Variants that were primary findings or actionable secondary findings were orthogonally confirmed regardless of the model’s predictions. Carrier status findings and pharmacogenomic findings were orthogonally confirmed when the model predicted the variant to be a false positive.

## RESULTS

### Variant Collection

HG001 (NA12878) was sequenced with three replicates and HG002 through HG005 were each sequenced once. The number of variants called across all samples was greater than 24 million true positive calls with 137 thousand false positive calls using the Dragen pipeline. Over 24 million true positive calls with 419 thousand false positive calls were found using the Sentieon/Strelka2 pipeline. Details of these counts by sample, variant type, and genotype along with a detailed description of the pipelines and RTG VCFeval invocations is available in the Supplemental Material.

### Model Evaluation

For our model selection and evaluation, we chose a minimum acceptable sensitivity of 0.99 (indicating 1/100 false calls are missed) with a target sensitivity of 0.995 (indicating 1/200 false calls are missed). Given these criteria, a number of models passed the evaluation process. The best model was chosen using the modified F1-score described above. The results for the final chosen models for all six variant-genotype combinations are shown in Table 1 for the Dragen pipeline and in Table 2 for the Sentieon/Strelka2 pipeline. Additional information for all final trained models at each evaluation sensitivity is available in the Supplemental Material.

**Table 1:** Summary of trained models for Dragen-based pipeline. For each variant-genotype combination, the following table reflects the best model for our criteria, the cross-validation (CV) mean and standard deviation for sensitivity and false positive rate (FPR), and final evaluation for sensitivity and FPR.

| Variant / Genotype | Best Model | CV Sensitivity | Final Sensitivity | CV FPR | Final FPR |
| --- | --- | --- | --- | --- | --- |
| SNV - Heterozygous | GradientBoosting | 0.9976+-0.0018 | 0.9958 | 0.1278+-0.0226 | 0.1220 |
| SNV - Homozygous | EasyEnsemble | 0.9994+-0.0014 | 0.9975 | 0.1725+-0.0207 | 0.1740 |
| SNV - Complex Het. | — | — | — | — | — |
| Indel - Heterozygous | GradientBoosting | 0.9962+-0.0026 | 0.9968 | 0.4311+-0.0335 | 0.4341 |
| Indel - Homozygous | GradientBoosting | 0.9978+-0.0027 | 0.9950 | 0.5565+-0.0416 | 0.5516 |
| Indel - Complex Het. | GradientBoosting | 0.9986+-0.0014 | 0.9960 | 0.5345+-0.0565 | 0.5422 |

**Table 2:** Summary of trained models for Sentieon/Strelka2-based pipeline. For each variant-genotype combination, the following table reflects the best model for our criteria, the cross-validation (CV) mean and standard deviation for sensitivity and false positive rate (FPR), and final evaluation for sensitivity and FPR. Note that no model passed our criteria for the complex heterozygous SNV dataset.

| Variant / Genotype | Best Model | CV Sensitivity | Final Sensitivity | CV FPR | Final FPR |
| --- | --- | --- | --- | --- | --- |
| SNV - Heterozygous | GradientBoosting | 0.9958+-0.0007 | 0.9952 | 0.0166+-0.0026 | 0.0167 |
| SNV - Homozygous | EasyEnsemble | 0.9987+-0.0032 | 0.9955 | 0.1543+-0.0566 | 0.1483 |
| SNV - Complex Het. | — | — | — | — | — |
| Indel - Heterozygous | GradientBoosting | 0.9958+-0.0011 | 0.9950 | 0.2040+-0.0328 | 0.2029 |
| Indel - Homozygous | GradientBoosting | 0.9968+-0.0015 | 0.9955 | 0.4235+-0.0398 | 0.4243 |
| Indel - Complex Het. | GradientBoosting | 0.9965+-0.0019 | 0.9955 | 0.6501+-0.0463 | 0.6495 |

Five of the six variant-genotype combinations had at least one model passing our criteria for both pipelines. The only failing combination was complex heterozygous SNVs (two non-reference alleles in trans at the same position), a failure that is likely due to the rarity of such events (45 false positive calls in the Dragen pipeline and 101 false positive calls in the Sentieon/Strelka2 pipeline across all seven samples). Models that were selected for use in clinical practice had a final sensitivity that was greater than or equal to our chosen target sensitivity of 0.9950. We tested a version of the models with very stringent criteria: minimum sensitivity of 0.9990 and a target sensitivity of 1.0000. Using these stringent models in conjunction with the Sentieon/Strelka2 pipeline resulted in a final false positive rate of 0.2802 for heterozygous SNVs and 0.6192 for heterozygous indels (see Supplemental Material for details).

### Clinical Evaluation

As we developed the models, we tracked Sanger confirmation results for cGS cases. The indication for testing was rare, undiagnosed disease. The first phase of the clinical evaluation was a retrospective analysis of recent cases for which Sanger confirmation results were available. We collected the orthogonal testing results for 232 variants from 26 cGS cases and compared them to the predictions from the Dragen-trained models. The results of this retrospective analysis are seen in Table 3. Only two variants in this dataset failed to confirm by orthogonal testing. Both were predicted to be false calls by the models. Of the 230 remaining true positive calls (i.e. confirmed by orthogonal testing), only 36 were incorrectly predicted to be false positives by the models. This indicates an observed false positive rate (FPR) of 0.1558 with observed model-specific FPRs ranging from 0.0294-0.2500.

**Table 3:**
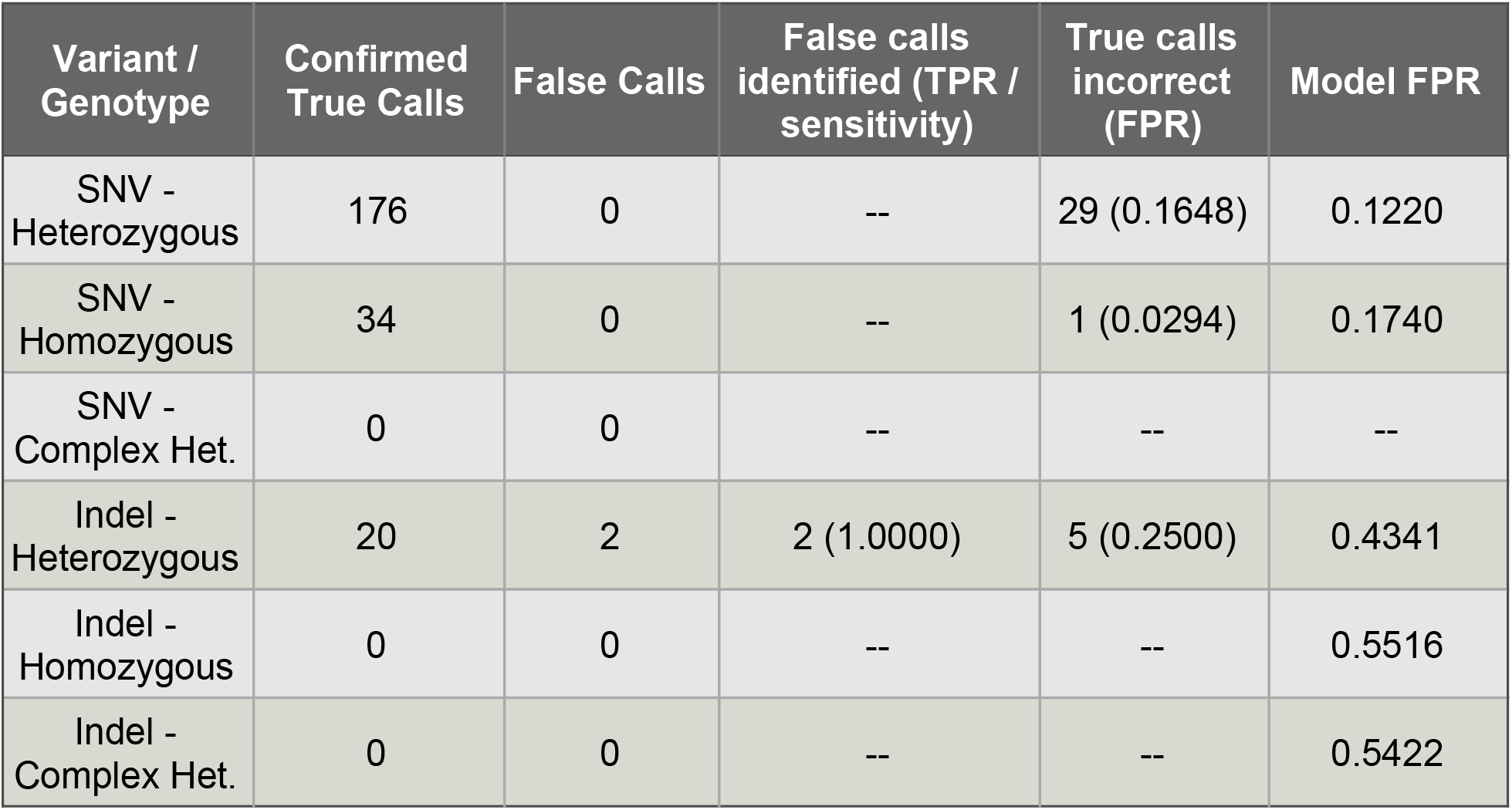
Summary of retrospective variant analysis. Here we report the total number of variants confirmed to be true positive or false positive calls, the number of false positive calls correctly identified (TPR / sensitivity), and the number of true calls incorrectly labeled as false calls (FPR). The model FPR (i.e., expected FPR) from the final evaluation is also provided here for comparison. Models used for this analysis were generated from the Dragen-based pipeline.

| Variant / Genotype | Confirmed True Calls | False Calls | False calls identified (TPR / sensitivity) | True calls incorrect (FPR) | Model FPR |
| --- | --- | --- | --- | --- | --- |
| SNV - Heterozygous | 176 | 0 | -- | 29 (0.1648) | 0.1220 |
| SNV - Homozygous | 34 | 0 | -- | 1 (0.0294) | 0.1740 |
| SNV - Complex Het. | 0 | 0 | -- | -- | -- |
| Indel - Heterozygous | 20 | 2 | 2 (1.0000) | 5 (0.2500) | 0.4341 |
| Indel - Homozygous | 0 | 0 | -- | -- | 0.5516 |
| Indel - Complex Het. | 0 | 0 | -- | -- | 0.5422 |

Following the development and evaluation noted above we employed the models in clinical practice to reduce the number of Sanger confirmations that were ordered in subsequent genome sequencing cases. As noted earlier, these models were only applied to non-actionable variants (carrier status findings and pharmacogenomic findings). Primary and actionable secondary variants continued to be sent for Sanger confirmation and were therefore excluded from this analysis. Additionally, every qualifying variant call that was predicted to be a true positive was manually reviewed using the Integrative Genomics Viewer^25^ as an additional review of the model’s prediction. We have applied the prediction algorithm to 252 non-actionable variants from 31 cGS cases gathered from the Dragen-based pipeline. Application of these models reduced the number of variants that had orthogonal confirmation by 216 (85.71%) overall, with an average reduction of 7.0 variants per sample. Sanger confirmation testing generally costs at least $100 (USD) per variant indicating an average cost savings of $696 per sample. Analysis of these results by variant type and genotype is shown in Table 4.

**Table 4:** Summary of prospective variant predictions. This table details the outcome of the use of the models in clinical cases. It shows the total number of variants that were predicted to be false positive or true positive in the clinical cases along with the percentage of variants that were not sent for orthogonal confirmation.

| Variant / Genotype | Predicted False Positive Calls | Predicted True Positive Calls | Orthogonal Order Reduction |
| --- | --- | --- | --- |
| SNV - Heterozygous | 29 | 164 | 84.97% |
| SNV - Homozygous | 1 | 34 | 97.14% |
| SNV - Complex Het. | 0 | 0 | -- |
| Indel - Heterozygous | 6 | 18 | 75.00% |
| Indel - Homozygous | 0 | 0 | -- |
| Indel - Complex Het. | 0 | 0 | -- |
| Overall | 36 | 216 | 85.71% |

## DISCUSSION

We developed a framework for training models to identify false positive variant calls from genome sequencing datasets. Our approach advances that of Lincoln et al.^10^ by using numerical values (rather than flags) as feature inputs to machine learning models, increasing the total number of true positive and false positive calls by using GIAB truth sets. This process obviates the need for a large set of orthogonal test results, a resource that is not available to most laboratories. In addition, the final models are tunable, allowing for laboratories to adjust the minimum and target sensitivities of the models to values that are relevant to a particular test application. Furthermore, the framework we developed can be used in conjunction with a variety of upstream pipelines as shown by the two different aligner-caller combinations used in this study. Custom models can also be developed to match upstream processes different from those used in this study such as different sequencing technologies, different secondary pipelines, and different major versions of the software used in a pipeline. The framework to develop custom models is available at https://github.com/HudsonAlpha/STEVE.

The model results in Tables 1 and 2 suggest that the upstream pipeline significantly influences the final performance of the models. For example, the final FPR rates (the fraction of true variants that would require orthogonal confirmation) are almost all lower with the Sentieon/Strelka2 pipeline. This suggests that while Sentieon/Strelka2 is generating more false positive calls (i.e. reduced precision compared to the Dragen pipeline), the features extracted from the VCF were better able to differentiate false positive calls from true positive calls compared to the features produced by the Dragen pipeline (see Supplemental Material). We anticipate that other pipelines will have similar variability in performance. Therefore we recommend building custom models for each upstream pipeline used.

Our results also suggest that there are differences between the full set of variants used in training and the set of variants that we are currently sending for orthogonal confirmation. For example, both heterozygous SNVs and indels had observed FPRs that are relatively close to the expected FPR of the Dragen model (see Table 3). In contrast, homozygous SNVs had an observed FPR (0.0294) much lower than the expected FPR from model evaluation (0.1740). While the numbers are too small to meet statistical significance, the results suggest that reported homozygous SNV variants are more likely to be predicted as true positives than homozygous SNV variants chosen at random. More data will be needed to assess this trend and determine whether similar trends occur for other variant types.

There are limitations and potential drawbacks to our approach. First, our approach is not trained on orthogonal results generated by Sanger sequencing. While generation of orthogonal results is an expensive process, it is possible that the feature distributions of variants that are clinically reported are different from those represented by the GIAB high confidence regions. For example, some clinically reportable variants are outside the GIAB high confidence regions. As a result, the models may not be trained to accurately predict the veracity of those variant calls. Additionally, many GIAB high-confidence variants are simply too common to be reported in a rare-disease clinical report. Those calls might dilute the performance of the models in some unexpected way. It is generally agreed that the interpretation of machine learning models is non-trivial and often referred to as a “black box”.15 While there are tools in place to aid in the interpretation of some models, they do not apply to all of the models we trained. As an example, the Supplement Materials detail some basic interpretation information referred to as “feature importances”, denoting which features are most influential in the models.^20^

We expect the performance of these models to improve as new GIAB truth sets and more replicates of existing GIAB truth sets are used in model training. Overall, these models show great promise to reduce orthogonal confirmation requirements while maintaining a low false positive rate of reported variants.

## ACKNOWLEDGEMENTS

We would like to thank the Smith Family Clinic and Children’s of Alabama for providing real patient samples and cases for evaluating the model performance in this paper. Sequencing was funded in part through the Hero Fund for Smith Family Clinic. We would like to thank all of the patients and families who allowed us to use their samples and data.

