## Supplemental document for "Reducing Sanger Confirmation Testing through False Positive Prediction Algorithms"

J. Matthew Holt et al.

April 21, 2020

### 1 Sample metadata

#### 1.1 General Sample Info

This section contains information regarding where samples were acquired from. This corresponds to the "Sample" label in Table 1.

1. NA12878 - female of European ancestry; purchased through `https://www.coriell.org/0/Sections/Search/Sample_Detail.aspx?Ref=NA12878&Product=DNA`
2. HG002-HG004 - son and parents of Eastern Europe Ashkenazi Jewish ancestry; purchased through `https://www-s.nist.gov/srmors/view_detail.cfm?srm=8392`
3. HG005 - male of Chinese ancestry; purchased through `https://www-s.nist.gov/srmors/view_detail.cfm?srm=8393`

#### 1.2 Training Samples

This section contains information regarding the specific samples used for analysis. This data is automatically pulled from a sample JSON file containing sample names, sample types (i.e. which GIAB sample), and how the sample was prepared. Table 1 contains the list of metadata as pulled from the JSON.

| Library | Sample | Preparation |
|---|---|---|
| SL362490 | NA12878 | Clinical PCR |
| SL362491 | NA12878 | Clinical PCR |
| SL362492 | NA12878 | Clinical PCR |
| SL409548 | HG002 | Clinical PCR |
| SL409549 | HG003 | Clinical PCR |
| SL409550 | HG004 | Clinical PCR |
| SL409551 | HG005 | Clinical PCR |

Table 1: This table contains metadata regarding each sequenced sample. The GIAB sample label and prep type are currently the two pieces of tracked metadata regarding each sample.

### 2 Genome Sequencing Pipelines

#### 2.1 Dragen Pipeline

Illumina's DRAGEN platform is a rapid genome analysis platform that performs both alignment and variant calling steps using hardware acceleration. The details of this platform can be found on Illumina's DRAGEN webpage.

##### 2.1.1 Integrated Command

Because the Dragen solution is fully integrated from FASTQ to gVCF, there is only one command we used to collect the final gVCFs. The final result of this step is the hard-filtered gVCF file (and corresponding index file) of the format `${sample}.hard-filtered.gvcf.gz`. That gVCF file is given to RTG VCFeval for variant evaluation.

```
dragen -f \
    -r /staging/reference/hg38/hg38.fa.k_21.f_16.m_149 \
    --fastq-list /staging/fastq/${sample}_fastqs/${sample}_list.csv \
    --bin_memory 60000000000 \
    --output-directory /staging/bam/ \
    --output-file-prefix ${sample} \
    --enable-duplicate-marking true \
    --enable-map-align-output true \
    --enable-variant-caller true \
    --vc-sample-name ${sample} \
    --vc-emit-ref-confidence GVCF \
    --dbsnp /staging/reference/hg38/dbsnp_146.hg38.vcf
```

#### 2.2 Sentieon / Strelka2 Pipeline

This pipeline uses a combination of Sentieon (more efficient implementation of BWA-mem) for alignment and Strelka2 for variant calling. The pipeline is implemented using a snakemake workflow, and relevent commands are presented in order below. All parameters referring to a reference genome are using the hg38 reference genome with ALT contigs.

##### 2.2.1 Sentieon paired-end alignment

The following command is used on each pair of FASTQ files for a sample. In brief, it performs the alignment process using sentieon, passes that into the post-alt alignment process derived from bwa-kit (this is recommended due to ALT contigs in the hg38 reference), and finally used the sention sorting function. The output of this command is a single, position-sorted BAM file that has been post-alt processed and the corresponding index file.

**Parameters:**

1. rgoptions - Read Group (RG) options for the particular flowcell/lane combination
2. reference - the filename for the reference genome (hg38 with all ALT contigs for our use case)
3. bwakit - directory containing a download of the bwa-kit post-ALT processing
4. tempParams - a temporary directory, can be removed without altering command outputs

```
sentieon \
    bwa mem -M \
    -R "{params.rgoptions}" \
    -t {threads} \
    -K 10000000 \
```

```
    {params.reference} \
    {input.fq1} {input.fq2} | \
{params.bwakit}/k8 \
    {params.bwakit}/bwa-postalt.js \
    {params.reference}.alt | \
sentieon util sort {params.tempParams} \
    --bam_compression 1 \
    -r {params.reference} \
    -o {output.bam} \
    -t {threads} \
    --sam2bam \
    -i -
```

###### 2.2.2 Sentieon deduplication

The following command will gather duplication statistics across *all* BAM files for a sample and then simultaneously remove duplicates while merging the BAM files together. The output of this step is a single BAM file containing all alignments for the sample and the corresponding index file.
**Parameters:**

1. sortedbams - this is a concatenation of `-i {BAM}` for each BAM file in the sample (i.e. each flowcell/lane BAM file generated in the previous step)

```
sentieon driver \
    -t {threads} \
    {params.sortedbams} \
    --algo LocusCollector \
    --fun score_info \
    {output.score} && \
sentieon driver {params.tempParams} \
    -t {threads} \
    {params.sortedbams} \
    --algo Dedup \
    --rmdup \
    --score_info {output.score} \
    --metrics {output.metrics} \
    --bam_compression 1 \
    {output.dedupbam}
```

###### 2.2.3 Sentieon Base Quality Recalibration

The following command will gather base quality score information for the de-duplicated sample BAM file and then perform base quality score recalibration (BQSR) on the BAM. The output of this step is a single BAM file containing the recalibrated mappings for the sample and the corresponding index file. This is the final BAM file for the sample.
**Parameters:**

1. reference - the filename for the reference genome (hg38 with all ALT contigs for our use case)
2. dbsnp - this is the dbSNP file gathered from this URL: `ftp://gsapubftp-anonymous@ftp.broadinstitute.org/bundle/hg38/dbsnp_146.hg38.vcf.gz`
3. mills - the is the Mills indel file gathered from this URL: `ftp://gsapubftp-anonymous@ftp.broadinstitute.org/bundle/hg38/Mills_and_1000G_gold_standard.indels.hg38.vcf.gz`
4. tempParams - a temporary directory, can be removed without altering command outputs

```
sentieon driver \
    -r {params.reference} \
    -t {threads} \
    -i {input.dedupbam} \
    --algo QualCal \
    -k {params.dbsnp} \
    -k {params.mills} \
    {output.recaltable} && \
sentieon driver {params.tempParams} \
    -r {params.reference} \
    -t {threads} \
    -i {input.dedupbam} \
    -q {output.recaltable} \
    --algo ReadWriter \
    {output.recalbam}
```

##### 2.2.4 Strelka2 Variant Calling

The following command will execute the Strelka2 workflow to perform variant calling. As an additional step, we annotate the final VCF file from Strelka2 with dbSNP identifiers (this is primarily for QC purposes in the pipeline). The final result of this step is a VCF file with dbSNP identifiers and the corresponding index file. This is the final VCF file that is provided as input to RTG VCFeval.
**Parameters:**

1. strelka - the path to the repo contain strelka2
2. reference - the filename for the reference genome (hg38 with all ALT contigs for our use case)
3. contigs - this is a restricted contig file (BED format) to reduce run time of Strelka2, see README file at `https://github.com/Illumina/strelka/blob/v2.9.x/docs/userGuide/README.md#improving-runtime-for-ref[illegible]` for the exact file and context behind usage
4. memGB - a memory limit for strelka2
5. bcftools - path to a bcftools executable for performing annotation; the version used was 1.10.2
6. dbsnp - this is the dbSNP file gathered from this URL: `ftp://gsapubftp-anonymous@ftp.broadinstitute.org/bundle/hg38/dbsnp_146.hg38.vcf.gz`

```
{params.strelka}/bin/configureStrelkaGermlineWorkflow.py \
    --bam {input.bam} \
    --referenceFasta {params.reference} \
    --callRegions {params.contigs} \
    --runDir {output.runDir} && \
{output.runDir}/runWorkflow.py \
    -m local \
    -j {threads} \
    -g {params.memGB} && \
{params.bcftools} annotate \
    -a {params.dbsnp} \
    -c ID \
    -O z \
    -o {output.vcf} \
    {output.runDir}/results/variants/variants.vcf.gz && \
tabix {output.vcf}
```

#### 2.3 RTG VCFeval Analysis

This tool was used to label variant calls as either true or false positives depending on presence or absence from the corresponding GIAB truth set. Note that these variants are limited to those found within the GIAB high-confidence regions (i.e. variants outside those regions are excluded).
**Parameters:**

1. truth - this is the VCF of variants published by GIAB representing a sample's truth set
2. bed - this is the high-confidence regions published by GIAB for the truth set; variants outside these regions are NOT evaluated
3. sdf - a file format required by RTG VCFeval (build from the hg38 reference)

```
rtg vcfeval \
    --all-records \
    -b {params.truth} \
    -c {input.vcf} \
    --bed-regions {params.bed} \
    -t {params.sdf} \
    -T {threads} \
    -o {output.rtgDir}
```

### 3 Model-Training Pipeline

This section contains details related to the methodology used for training all models. Note: an identical process is used for each pipeline, allowing for different configuration of inputs depending on the upstream pipeline.

#### 3.1 Feature Extraction

While an identical process is used for each pipeline, the features from each pipeline are configurable using a combination of JSON and hard-coded Python3 (when complex features are involved). Features must be numerical values when given to the models, so some transformations are necessary from the raw VCF specified values.

In the file `{REPO}/scripts/model_metrics.json`, there are a list of features defined for different upstream callers. When a feature is practically copied from a VCF file, we try to denote it below with the corresponding VCF tag. Here is a brief description of the sub-types and features (Note: not all types are used in each pipeline):

1. "CALL" - These features are generally tied to a genotype call (i.e. sample-specific)
   (a) AD0 - the allele depth (AD) for the first allele in the genotype (e.g. if GT=0/1, this is the depth of the reference allele)
   (b) AD1 - the allele depth (AD) for the second allele in the genotype (e.g. if GT=0/1, this is the depth of the first alternate allele)
   (c) ADO - the total allele depth (AD) for any alleles that are not present in the genotype call
   (d) AF0 - the allele frequency for the first allele in the genotype (e.g. if GT=0/1 and AD=10,30 then this value is 0.25)
   (e) AF1 - the allele frequency for the second allele in the genotype (e.g. if GT=0/1 and AD=10,30 then this value is 0.75)
   (f) AFO - the total allele frequency for any alleles that are not present in the genotype call
   (g) GT - the genotype field (GT) transformed into a single numerical value
   (h) DP - the depth field (DP)
   (i) GQ - the genotype quality (GQ) field
   (j) DPI - the indel read depth (DPI)
   (k) GQX - empirically calibrated genotype quality score (GQX)
   (l) DPF - basecalls filtered prior to genotyping (DPF)
   (m) SB - sample site strand bias (SB)
2. "INFO" - These features are generally tied to a variant site and may represent aggregate quality statistics in multi-sample VCF files (i.e. variant-specific metrics)
   (a) DB - represents dbSNP membership (DB)
   (b) FractionInformativeReads - fraction of informative reads out of the total reads (FractionInformativeReads)
   (c) FS - Phred-scaled Fisher's Exact Test for strand bias (FS)
   (d) MQ - mapping quality (MQ)
   (e) MQRankSum - rank sum test for mapping qualities (MQRankSum)
   (f) QD - variant confidence by depth (QD)
   (g) R2_5P_bias - score based on mate bias and distance from 5-prime end (R2_5P_bias)
   (h) ReadPosRankSum - measure of position bias (ReadPosRankSum)

(i) SOR - measure of strand bias using contingency table (SOR)
(j) SNVHPOL - SNV context homopolymer length (SNVHPOL)

3. "MUNGED" - These features are generally calculated from information present in the VCF files that does not cleanly fall into either the INFO or CALL feature types
   (a) DP_DP - ratio of call depth over total variant depth (generally 1.0 for single-sample VCFs)
   (b) QUAL - the quality value in the VCF (QUAL)
   (c) NEARBY - the number of non-reference variant calls near the current variant ($\pm$20bp)
   (d) FILTER - the number of non-PASS filter values in the FILTER field of the VCF
   (e) ID - set to True (i.e. 1) if the ID field is not empty, otherwise False (i.e. 0)

#### 3.2 Model Hyperparameters

During cross-validation, the models are given a selection of hyperparameters (i.e. parameters that define how the models are built) to choose from to identify the "best" combination of hyperparameters for the particular dataset. We selected a handful of hyperparameters based on the recommendations provided by `sklearn`, `imblearn`, and/or the corresponding literature for the models. We then applied `sklearn`'s `GridSearchSV` method which systematically tests every possible combination of hyperparameters of those provided. Only the best hyperparameters are then used for the final training and testing.

Table 2 reproduces the list of hyperparameters that were initially tested. Note that this list is not exhaustive, but is intended to represent the most impactful hyperparameters. Additionally, this list of hyperparameters is statically entered into this document, but it is subject to change with new versions and is best found embedded within the source code in file `{REPO}/scripts/TrainModels.py`.

| Model | Hyperparameter | Search Space |
|---|---|---|
| RandomForestClassifier (sklearn) | random_state | [0] |
| | class_weight | ['balanced'] |
| | n_estimators | [100, 200] |
| | max_depth | [3, 4] |
| | min_samples_split | [2] |
| | max_features | ['sqrt'] |
| AdaBoostClassifier (sklearn) | random_state | [0] |
| | base_estimator | [DecisionTreeClassifier(max_depth=2)] |
| | n_estimators | [100, 200] |
| | learning_rate | [0.01, 0.1, 1.0] |
| | algorithm | ['SAMME', 'SAMME.R'] |
| GradientBoostingClassifier (sklearn) | random_state | [0] |
| | n_estimators | [100, 200] |
| | max_depth | [3, 4] |
| | learning_rate | [0.05, 0.1, 0.2] |
| | loss | ['deviance', 'exponential'] |
| | max_features | ['sqrt'] |
| EasyEnsembleClassifier (imblearn) | random_state | [0] |
| | n_estimators | [10, 20, 30, 40, 50] |

Table 2: Hyperparameters tested in the initial version of the training pipeline.

#### 3.3 Clinical Model Selection Formula

After full training, the models are evaluated on the unseen test dataset. Any candidate models are required to pass a cross-validation sensitivity requirement and final sensitivity requirement (see main document for details). We then use the following methodology to select the "best" candidate model that will ultimately be used clinically. Note that this process is used for each variant/genotype combination, culminating in up to six final models (one per combo).

1. Let $S_m = 0.99$ be the minimum acceptable sensitivity and $S_t = 0.995$ be the target sensitivity for the models.
2. For each candidate model, let $S$ be the final sensitivity and $F$ be the final false positive rate (FPR) for the model.
3. Calculate the scaled sensitivity score, that is at most 1.0 (representing a model reaching the target sensitivity): $S_s = min(1.0, \frac{S-S_m}{S_t-S_m})$
4. Calculate the specificity (true negative rate), $T = 1.0 - F$, such that higher values indicate fewer true variant calls being incorrectly sent for confirmation.
5. Calculate the modified F1 score: $F = harmonic\_mean(S_s, T)$
6. Of the remaining models, select the model with the highest F1 score, $F$, for use clinically.

| Sample | True Positives | False Positives | Sensitivity | Precision | F-measure |
|---|---|---|---|---|---|
| SL362490 | 3,526,846 | 17,164 | 0.9955 | 0.9952 | 0.9954 |
| SL362491 | 3,527,734 | 16,757 | 0.9958 | 0.9953 | 0.9955 |
| SL362492 | 3,530,085 | 17,184 | 0.9965 | 0.9952 | 0.9958 |
| SL409548 | 3,485,975 | 22,200 | 0.9945 | 0.9937 | 0.9941 |
| SL409549 | 3,336,817 | 20,287 | 0.9948 | 0.9940 | 0.9944 |
| SL409550 | 3,368,531 | 30,167 | 0.9937 | 0.9911 | 0.9924 |
| SL409551 | 3,298,988 | 13,388 | 0.9971 | 0.9960 | 0.9965 |
| Mean±Stdev | 3,439,282±93,430 | 19,592±5,033 | 0.9954±0.0011 | 0.9944±0.0015 | 0.9949±0.0013 |

Table 3: Summary metrics from RTG VCFeval for aligner "dragen-07.011.352.3.2.8b" and variant caller "dragen-07.011.352.3.2.8b".

### 4 Results for dragen-07.011.352.3.2.8b/dragen-07.011.352.3.2.8b

The following sections denote results that are specific the the pipeline consisting of aligner "dragen-07.011.352.3.2.8b" and variant caller "dragen-07.011.352.3.2.8b".

#### 4.1 RTG VCFeval Results

The following sections contain results as reported by `rtg vcfeval`. For information on how `rtg vcfeval` was invoked, refer to Section 2.3.

##### 4.1.1 Pipeline Performance

Table 3 contains the results from the RTG VCFeval `summary.txt` file that primarily contains summary information regarding the evaluated VCF file. We copied the results from this summary (unfiltered "None" row) and calculated summary mean and standard deviation as well.

Sensitivity is the fraction of annotated true positives that were correctly identified by the pipeline, precision is the fraction of called variants that were part of the truth set, and F-measure is the harmonic mean of sensitivy and precision. A perfect caller would have 1.0000 for all scores.

##### 4.1.2 Variant Counts

Table 4 contains a summary of the number of false and true positive variant calls after stratifying the results by variant type and genotype.

#### 4.2 Model Results

The following sections contain results specific to the final trained models.

##### 4.2.1 Selected Models

Table 5 contains the selected models for aligner "dragen-07.011.352.3.2.8b" and caller "dragen-07.011.352.3.2.8b" given the minimum sensitivity, $S_m = 0.99$, and the target sensitivity, $S_t = 0.995$.

##### 4.2.2 Strict Models

Table 6 contains the strict models for aligner "dragen-07.011.352.3.2.8b" and caller "dragen-07.011.352.3.2.8b" given the minimum sensitivity, $S_m = 0.999$, and the target sensitivity, $S_t = 1.0$. These models are labeled strict due to very high requirements, and the majority of models at different evaluation sensitivities fail to pass these criteria. As a result, many variant/genotype combinations have no passing models or have models that are not practically useful (e.g. an FPR of 99%).

| Sample | RTG Result | SNV-HET | SNV-HOM | SNV-HE2 | INDEL-HET | INDEL-HOM | INDEL-HE2 | Total Calls |
|---|---|---|---|---|---|---|---|---|
| SL362490 | FP | 3,133 | 543 | 5 | 9,351 | 1,913 | 2,217 | 17,162 |
| SL362491 | FP | 2,997 | 521 | 3 | 9,224 | 1,806 | 2,205 | 16,756 |
| SL362492 | FP | 3,372 | 484 | 8 | 9,734 | 1,260 | 2,325 | 17,183 |
| SL409548 | FP | 5,527 | 956 | 8 | 11,589 | 1,536 | 2,584 | 22,200 |
| SL409549 | FP | 4,583 | 721 | 8 | 10,933 | 1,424 | 2,617 | 20,286 |
| SL409550 | FP | 6,018 | 770 | 5 | 16,898 | 2,156 | 4,319 | 30,166 |
| SL409551 | FP | 4,553 | 712 | 8 | 5,870 | 1,208 | 1,036 | 13,387 |
| Total | FP | 30,183 | 4,707 | 45 | 73,599 | 11,303 | 17,303 | 137,140 |
| SL362490 | TP | 1,845,739 | 1,192,614 | 838 | 285,389 | 170,848 | 31,416 | 3,526,844 |
| SL362491 | TP | 1,845,930 | 1,192,707 | 835 | 285,750 | 170,952 | 31,559 | 3,527,733 |
| SL362492 | TP | 1,846,038 | 1,192,575 | 833 | 287,380 | 171,015 | 32,243 | 3,530,084 |
| SL409548 | TP | 1,860,569 | 1,160,289 | 889 | 275,934 | 160,381 | 27,909 | 3,485,971 |
| SL409549 | TP | 1,742,984 | 1,132,667 | 824 | 271,048 | 162,620 | 26,674 | 3,336,817 |
| SL409550 | TP | 1,780,448 | 1,122,038 | 808 | 276,683 | 161,507 | 27,047 | 3,368,531 |
| SL409551 | TP | 1,646,670 | 1,253,290 | 792 | 224,184 | 153,317 | 20,734 | 3,298,987 |
| Total | TP | 12,568,378 | 8,246,180 | 5,819 | 1,906,368 | 1,150,640 | 197,582 | 24,074,967 |

Table 4: This table shows the number of false and true positive variants calls as reported by `rtg vcfeval` for the aligner dragen-07.011.352.3.2.8b and variant caller dragen-07.011.352.3.2.8b. The variants are further divided by variant type (SNV or INDEL) and genotype (HET=heterozygous, HOM=homozygous, HE2=complex heterozygous). The "total" label refers to the sum of all samples for the corresponding "RTG Result" type.

| Variant type | Best Model | Evaluation Sensitivity | CV Sensitivity | Final Sensitivity | CV FPR | Final FPR |
|---|---|---|---|---|---|---|
| SNV-HET | GradientBoosting | 0.998 | 0.9976+-0.0018 | 0.9958 | 0.1278+-0.0226 | 0.1220 |
| SNV-HOM | EasyEnsemble | 0.9999 | 0.9994+-0.0014 | 0.9975 | 0.1725+-0.0207 | 0.1740 |
| SNV-HE2 | None | None | – | – | – | – |
| INDEL-HET | GradientBoosting | 0.997 | 0.9962+-0.0026 | 0.9968 | 0.4311+-0.0335 | 0.4341 |
| INDEL-HOM | GradientBoosting | 0.998 | 0.9978+-0.0027 | 0.9950 | 0.5565+-0.0416 | 0.5516 |
| INDEL-HE2 | GradientBoosting | 0.999 | 0.9986+-0.0014 | 0.9960 | 0.5345+-0.0565 | 0.5422 |

Table 5: Selected models for aligner "dragen-07.011.352.3.2.8b", caller "dragen-07.011.352.3.2.8b", $S_m = 0.99$, $S_t = 0.995$. If no model passed the criteria, then the "Best Model" field will be "None". Evaluation sensitivity is the training sensitivity that was used to gather results for the remaining fields in testing. Results prefaced with "CV" represent the test results during cross-validation. Similarly, results prefaced with "Final" represent the results on the held-out testing set during final evaluation. Note that we required the models to have sensitivity requirements based on both the CV and Final results. In contrast, FPR is not bound by any requirements, but is instead representative of the expected fraction of orthogonal confirmations required if the model is used.

| Variant type | Best Model | Evaluation Sensitivity | CV Sensitivity | Final Sensitivity | CV FPR | Final FPR |
|---|---|---|---|---|---|---|
| SNV-HET | EasyEnsemble | 1.0 | 0.9999+-0.0002 | 0.9999 | 0.8805+-0.0239 | 0.8885 |
| SNV-HOM | None | None | – | – | – | – |
| SNV-HE2 | None | None | – | – | – | – |
| INDEL-HET | GradientBoosting | 0.9999 | 0.9999+-0.0002 | 0.9999 | 0.7957+-0.0536 | 0.8144 |
| INDEL-HOM | AdaBoost | 0.9999 | 0.9998+-0.0004 | 0.9995 | 0.8404+-0.0329 | 0.8414 |
| INDEL-HE2 | RandomForest | 0.9999 | 0.9999+-0.0003 | 0.9998 | 0.9767+-0.0228 | 0.9867 |

Table 6: Strict models for aligner "dragen-07.011.352.3.2.8b", caller "dragen-07.011.352.3.2.8b", $S_m = 0.999$, $S_t = 1.0$. If no model passed the criteria, then the "Best Model" field will be "None". Evaluation sensitivity is the training sensitivity that was used to gather results for the remaining fields in testing. Results prefaced with "CV" represent the test results during cross-validation. Similarly, results prefaced with "Final" represent the results on the held-out testing set during final evaluation. Note that we required the models to have sensitivity requirements based on both the CV and Final results. In contrast, FPR is not bound by any requirements, but is instead representative of the expected fraction of orthogonal confirmations required if the model is used.

###### 4.2.3 Feature Importances

Table 7 contains results regarding feature importances according to the models. These were gathered using the `eli5` package and the `ExtractELI5Results.py` script from this repo. Feature importances may be missing due to any of the following reasons:

1. ELI5 interpretation was not run correctly - This could be because the `ExtractELI5Results.py` script has not been executed or the outputs are not in the expected location.
2. The model failed to pass our base clinical criteria - We restricted the outputs to only include models that met the minimum sensitivity requirement as defined in the "Selected Models" section above.
3. The model is not interpretable by `eli5` - Not all models provide feature importance measures through eli5 so these results are excluded

###### 4.2.4 Model for SNV-HET

Figure 1 contains the receiver-operator curves (ROC) for the final trained models for aligner "dragen-07.011.352.3.2.8b", caller "dragen-07.011.352.3.2.8b", variant type "SNV", and genotype "HET".

| **Feature** | SNV-HET | SNV-HOM | SNV-HE2 | INDEL-HET | INDEL-HOM | INDEL-HE2 | Cumulative |
|---|---|---|---|---|---|---|---|
| CALL-GQ | 0.3293 | – | – | 0.4679 | 0.2290 | 0.6180 | 1.6442 |
| MUNGED-FILTER | 0.1462 | – | – | 0.2199 | 0.0000 | 0.0002 | 0.3663 |
| CALL-AD0 | 0.0265 | – | – | 0.0077 | 0.2243 | 0.0280 | 0.2866 |
| CALL-AF1 | 0.0756 | – | – | 0.0968 | 0.0201 | 0.0811 | 0.2736 |
| MUNGED-QUAL | 0.0444 | – | – | 0.0759 | 0.0686 | 0.0720 | 0.2609 |
| INFO-DB | 0.1511 | – | – | 0.0338 | 0.0275 | 0.0184 | 0.2308 |
| CALL-DP | 0.0111 | – | – | 0.0034 | 0.1712 | 0.0185 | 0.2041 |
| CALL-AD1 | 0.0162 | – | – | 0.0239 | 0.1164 | 0.0461 | 0.2025 |
| MUNGED-NEARBY | 0.0688 | – | – | 0.0156 | 0.0444 | 0.0079 | 0.1367 |
| CALL-AF0 | 0.0173 | – | – | 0.0201 | 0.0234 | 0.0660 | 0.1268 |
| INFO-FractionInformativeReads | 0.0236 | – | – | 0.0062 | 0.0351 | 0.0069 | 0.0718 |
| INFO-MQ | 0.0374 | – | – | 0.0071 | 0.0148 | 0.0074 | 0.0667 |
| INFO-MQRankSum | 0.0224 | – | – | 0.0128 | 0.0069 | 0.0142 | 0.0563 |
| MUNGED-DP_DP | 0.0207 | – | – | 0.0050 | 0.0108 | 0.0071 | 0.0436 |
| INFO-ReadPosRankSum | 0.0077 | – | – | 0.0039 | 0.0047 | 0.0056 | 0.0218 |
| INFO-R2_5P_bias | 0.0018 | – | – | 0.0002 | 0.0028 | 0.0024 | 0.0072 |
| INFO-QD | 0.0000 | – | – | 0.0000 | 0.0000 | 0.0000 | 0.0000 |
| INFO-FS | 0.0000 | – | – | 0.0000 | 0.0000 | 0.0000 | 0.0000 |
| INFO-SOR | 0.0000 | – | – | 0.0000 | 0.0000 | 0.0000 | 0.0000 |

Table 7: This table shows the feature importances results for aligner “dragen-07.011.352.3.2.8b” and caller “dragen-07.011.352.3.2.8b”. Importances are broken down by category with a cumulative sum at the end. Note that some results may be missing if the pipeline was run incorrectly or the models are not interpretable through `eli5`.

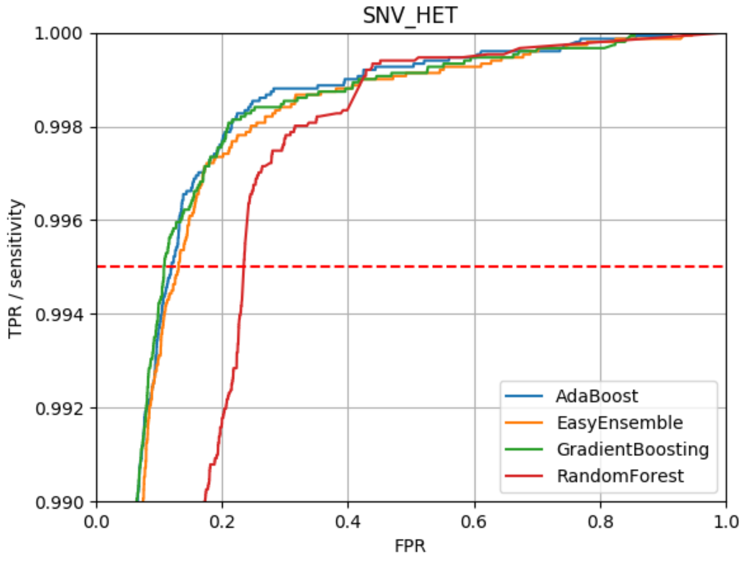

Figure 1: ROC curve for aligner “dragen-07.011.352.3.2.8b”, caller “dragen-07.011.352.3.2.8b”, variant type “SNV”, and genotype “HET”. Note that these curves are zoomed in to focus on only the region greater than the minimum clinical sensitivity (0.99).

###### 4.2.5 Model for SNV-HOM

Figure 2 contains the receiver-operator curves (ROC) for the final trained models for aligner “dragen-07.011.352.3.2.8b”, caller “dragen-07.011.352.3.2.8b”, variant type “SNV”, and genotype “HOM”.

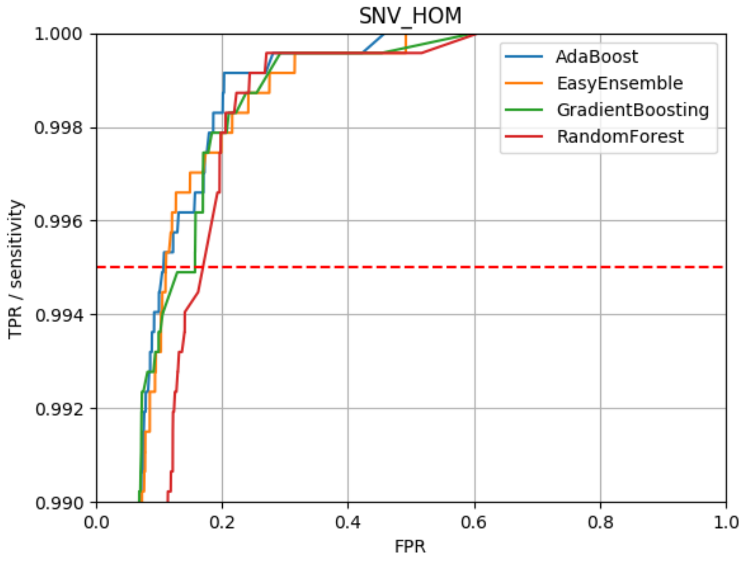

Figure 2: ROC curve for aligner “dragen-07.011.352.3.2.8b”, caller “dragen-07.011.352.3.2.8b”, variant type “SNV”, and genotype “HOM”. Note that these curves are zoomed in to focus on only the region greater than the minimum clinical sensitivity (0.99).

###### 4.2.6 Model for SNV-HE2

Figure 3 contains the receiver-operator curves (ROC) for the final trained models for aligner "dragen-07.011.352.3.2.8b", caller "dragen-07.011.352.3.2.8b", variant type "SNV", and genotype "HE2".

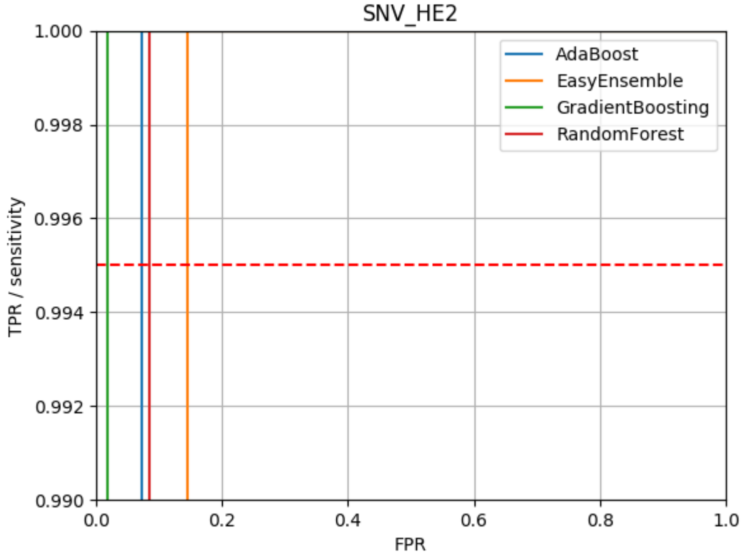

Figure 3: ROC curve for aligner "dragen-07.011.352.3.2.8b", caller "dragen-07.011.352.3.2.8b", variant type "SNV", and genotype "HE2". Note that these curves are zoomed in to focus on only the region greater than the minimum clinical sensitivity (0.99).

###### 4.2.7 Model for INDEL-HET

Figure 4 contains the receiver-operator curves (ROC) for the final trained models for aligner "dragen-07.011.352.3.2.8b", caller "dragen-07.011.352.3.2.8b", variant type "INDEL", and genotype "HET".

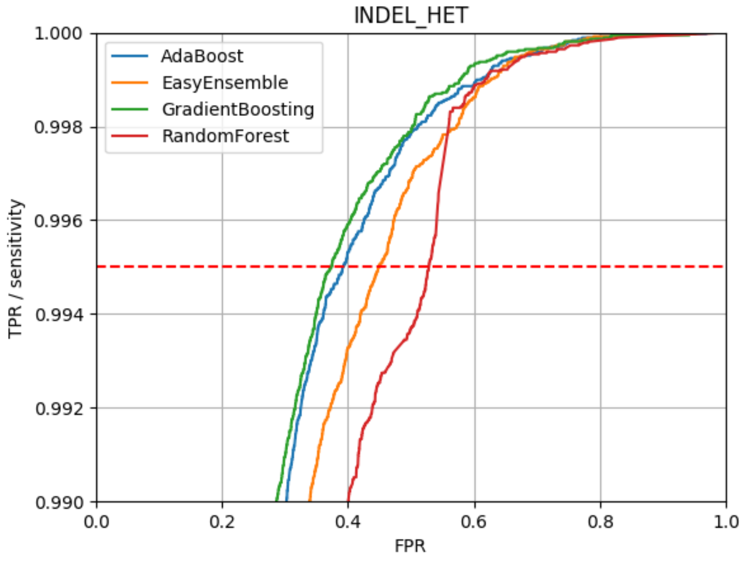

Figure 4: ROC curve for aligner "dragen-07.011.352.3.2.8b", caller "dragen-07.011.352.3.2.8b", variant type "INDEL", and genotype "HET". Note that these curves are zoomed in to focus on only the region greater than the minimum clinical sensitivity (0.99).

###### 4.2.8 Model for INDEL-HOM

Figure 5 contains the receiver-operator curves (ROC) for the final trained models for aligner "dragen-07.011.352.3.2.8b", caller "dragen-07.011.352.3.2.8b", variant type "INDEL", and genotype "HOM".

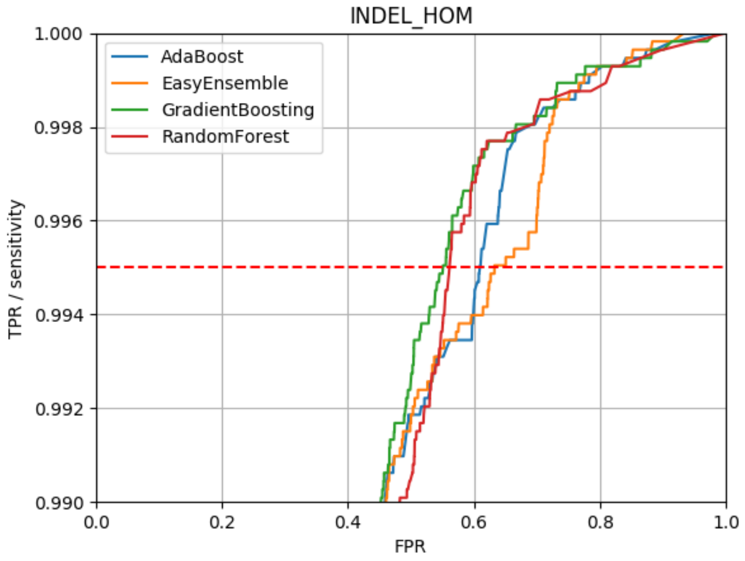

Figure 5: ROC curve for aligner "dragen-07.011.352.3.2.8b", caller "dragen-07.011.352.3.2.8b", variant type "INDEL", and genotype "HOM". Note that these curves are zoomed in to focus on only the region greater than the minimum clinical sensitivity (0.99).

###### 4.2.9 Model for INDEL-HE2

Figure 6 contains the receiver-operator curves (ROC) for the final trained models for aligner "dragen-07.011.352.3.2.8b", caller "dragen-07.011.352.3.2.8b", variant type "INDEL", and genotype "HE2".

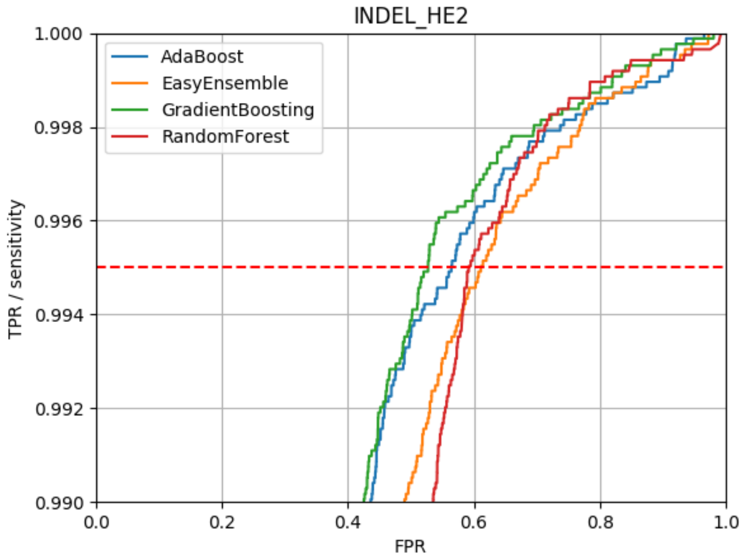

Figure 6: ROC curve for aligner "dragen-07.011.352.3.2.8b", caller "dragen-07.011.352.3.2.8b", variant type "INDEL", and genotype "HE2". Note that these curves are zoomed in to focus on only the region greater than the minimum clinical sensitivity (0.99).

| Sample | True Positives | False Positives | Sensitivity | Precision | F-measure |
|---|---|---|---|---|---|
| SL362490 | 3,525,149 | 54,392 | 0.9948 | 0.9848 | 0.9898 |
| SL362491 | 3,526,218 | 51,432 | 0.9951 | 0.9856 | 0.9903 |
| SL362492 | 3,530,305 | 48,127 | 0.9962 | 0.9866 | 0.9914 |
| SL409548 | 3,485,151 | 66,280 | 0.9940 | 0.9813 | 0.9876 |
| SL409549 | 3,336,429 | 61,613 | 0.9944 | 0.9819 | 0.9881 |
| SL409550 | 3,366,507 | 88,197 | 0.9928 | 0.9745 | 0.9836 |
| SL409551 | 3,297,011 | 49,890 | 0.9963 | 0.9851 | 0.9907 |
| Mean±Stdev | 3,438,110±93,676 | 59,990±13,012 | 0.9948±0.0011 | 0.9828±0.0038 | 0.9888±0.0025 |

Table 8: Summary metrics from RTG VCFeval for aligner "sentieon-201808.07" and variant caller "strelka-2.9.10".

### 5 Results for sentieon-201808.07/strelka-2.9.10

The following sections denote results that are specific the the pipeline consisting of aligner "sentieon-201808.07" and variant caller "strelka-2.9.10".

#### 5.1 RTG VCFeval Results

The following sections contain results as reported by `rtg vcfeval`. For information on how `rtg vcfeval` was invoked, refer to Section 2.3.

##### 5.1.1 Pipeline Performance

Table 8 contains the results from the RTG VCFeval `summary.txt` file that primarily contains summary information regarding the evaluated VCF file. We copied the results from this summary (unfiltered "None" row) and calculated summary mean and standard deviation as well.

Sensitivity is the fraction of annotated true positives that were correctly identified by the pipeline, precision is the fraction of called variants that were part of the truth set, and F-measure is the harmonic mean of sensitivy and precision. A perfect caller would have 1.0000 for all scores.

##### 5.1.2 Variant Counts

Table 9 contains a summary of the number of false and true positive variant calls after stratifying the results by variant type and genotype.

#### 5.2 Model Results

The following sections contain results specific to the final trained models.

##### 5.2.1 Selected Models

Table 10 contains the selected models for aligner "sentieon-201808.07" and caller "strelka-2.9.10" given the minimum sensitivity, $S_m = 0.99$, and the target sensitivity, $S_t = 0.995$.

##### 5.2.2 Strict Models

Table 11 contains the strict models for aligner "sentieon-201808.07" and caller "strelka-2.9.10" given the minimum sensitivity, $S_m = 0.999$, and the target sensitivity, $S_t = 1.0$. These models are labeled strict due to very high requirements, and the majority of models at different evaluation sensitivities fail to pass these criteria. As a result, many variant/genotype combinations have no passing models or have models that are not practically useful (e.g. an FPR of 99%).

| Sample | RTG Result | SNV-HET | SNV-HOM | SNV-HE2 | INDEL-HET | INDEL-HOM | INDEL-HE2 | Total Calls |
|---|---|---|---|---|---|---|---|---|
| SL362490 | FP | 19,475 | 316 | 13 | 28,467 | 4,681 | 1,369 | 54,321 |
| SL362491 | FP | 16,960 | 261 | 9 | 28,009 | 4,781 | 1,345 | 51,365 |
| SL362492 | FP | 13,941 | 200 | 17 | 28,423 | 4,117 | 1,370 | 48,068 |
| SL409548 | FP | 24,560 | 434 | 16 | 35,297 | 4,239 | 1,635 | 66,181 |
| SL409549 | FP | 19,935 | 396 | 11 | 35,550 | 3,966 | 1,680 | 61,538 |
| SL409550 | FP | 37,516 | 343 | 25 | 41,365 | 6,695 | 2,155 | 88,099 |
| SL409551 | FP | 27,772 | 278 | 10 | 18,150 | 2,931 | 683 | 49,824 |
| Total | FP | 160,159 | 2,228 | 101 | 215,261 | 31,410 | 10,237 | 419,396 |
| SL362490 | TP | 1,848,561 | 1,192,431 | 837 | 283,154 | 171,678 | 28,315 | 3,524,976 |
| SL362491 | TP | 1,848,751 | 1,192,465 | 840 | 283,699 | 171,887 | 28,395 | 3,526,037 |
| SL362492 | TP | 1,849,072 | 1,192,443 | 841 | 285,800 | 172,349 | 29,655 | 3,530,160 |
| SL409548 | TP | 1,863,668 | 1,159,651 | 890 | 273,971 | 161,656 | 25,104 | 3,484,940 |
| SL409549 | TP | 1,745,901 | 1,132,245 | 825 | 269,263 | 163,809 | 24,177 | 3,336,220 |
| SL409550 | TP | 1,783,155 | 1,121,260 | 812 | 273,794 | 163,741 | 23,550 | 3,366,312 |
| SL409551 | TP | 1,648,579 | 1,252,589 | 793 | 222,881 | 153,310 | 18,756 | 3,296,908 |
| Total | TP | 12,587,687 | 8,243,084 | 5,838 | 1,892,562 | 1,158,430 | 177,952 | 24,065,553 |

Table 9: This table shows the number of false and true positive variants calls as reported by `rtg vcfeval` for the aligner sentieon-201808.07 and variant caller strelka-2.9.10. The variants are further divided by variant type (SNV or INDEL) and genotype (HET=heterozygous, HOM=homozygous, HE2=complex heterozygous). The "total" label refers to the sum of all samples for the corresponding "RTG Result" type.

| Variant type | Best Model | Evaluation Sensitivity | CV Sensitivity | Final Sensitivity | CV FPR | Final FPR |
|---|---|---|---|---|---|---|
| SNV-HET | GradientBoosting | 0.996 | 0.9958+-0.0007 | 0.9952 | 0.0166+-0.0026 | 0.0167 |
| SNV-HOM | EasyEnsemble | 0.998 | 0.9987+-0.0032 | 0.9955 | 0.1543+-0.0566 | 0.1483 |
| SNV-HE2 | None | None | – | – | – | – |
| INDEL-HET | GradientBoosting | 0.996 | 0.9958+-0.0011 | 0.9950 | 0.2040+-0.0328 | 0.2029 |
| INDEL-HOM | GradientBoosting | 0.997 | 0.9968+-0.0015 | 0.9955 | 0.4235+-0.0398 | 0.4243 |
| INDEL-HE2 | GradientBoosting | 0.997 | 0.9965+-0.0019 | 0.9955 | 0.6501+-0.0463 | 0.6495 |

Table 10: Selected models for aligner "sentieon-201808.07", caller "strelka-2.9.10", $S_m = 0.99$, $S_t = 0.995$. If no model passed the criteria, then the "Best Model" field will be "None". Evaluation sensitivity is the training sensitivity that was used to gather results for the remaining fields in testing. Results prefaced with "CV" represent the test results during cross-validation. Similarly, results prefaced with "Final" represent the results on the held-out testing set during final evaluation. Note that we required the models to have sensitivity requirements based on both the CV and Final results. In contrast, FPR is not bound by any requirements, but is instead representative of the expected fraction of orthogonal confirmations required if the model is used.

| Variant type | Best Model | Evaluation Sensitivity | CV Sensitivity | Final Sensitivity | CV FPR | Final FPR |
|---|---|---|---|---|---|---|
| SNV-HET | GradientBoosting | 0.9999 | 0.9999+-0.0002 | 0.9997 | 0.2863+-0.0344 | 0.2802 |
| SNV-HOM | RandomForest | 0.999 | 1.0000+-0.0000 | 1.0000 | 0.9973+-0.0021 | 0.9972 |
| SNV-HE2 | None | None | – | – | – | – |
| INDEL-HET | GradientBoosting | 0.9999 | 0.9999+-0.0002 | 0.9997 | 0.6160+-0.0492 | 0.6192 |
| INDEL-HOM | EasyEnsemble | 0.9999 | 0.9999+-0.0004 | 0.9997 | 0.9258+-0.0200 | 0.9296 |
| INDEL-HE2 | None | None | – | – | – | – |

Table 11: Strict models for aligner “sentieon-201808.07”, caller “strelka-2.9.10”, $S_m = 0.999$, $S_t = 1.0$. If no model passed the criteria, then the “Best Model” field will be “None”. Evaluation sensitivity is the training sensitivity that was used to gather results for the remaining fields in testing. Results prefaced with “CV” represent the test results during cross-validation. Similarly, results prefaced with “Final” represent the results on the held-out testing set during final evaluation. Note that we required the models to have sensitivity requirements based on both the CV and Final results. In contrast, FPR is not bound by any requirements, but is instead representative of the expected fraction of orthogonal confirmations required if the model is used.

###### 5.2.3 Feature Importances

Table 12 contains results regarding feature importances according to the models. These were gathered using the `eli5` package and the `ExtractELI5Results.py` script from this repo. Feature importances may be missing due to any of the following reasons:

1. ELI5 interpretation was not run correctly - This could be because the `ExtractELI5Results.py` script has not been executed or the outputs are not in the expected location.
2. The model failed to pass our base clinical criteria - We restricted the outputs to only include models that met the minimum sensitivity requirement as defined in the “Selected Models” section above.
3. The model is not interpretable by `eli5` - Not all models provide feature importance measures through eli5 so these results are excluded

###### 5.2.4 Model for SNV-HET

Figure 7 contains the receiver-operator curves (ROC) for the final trained models for aligner “sentieon-201808.07”, caller “strelka-2.9.10”, variant type “SNV”, and genotype “HET”.

| Feature | SNV-HET | SNV-HOM | SNV-HE2 | INDEL-HET | INDEL-HOM | INDEL-HE2 | Cumulative |
|---|---|---|---|---|---|---|---|
| CALL-GQX | 0.1562 | – | – | 0.3838 | 0.4500 | 0.3621 | 1.3521 |
| MUNGED-QUAL | 0.0122 | – | – | 0.4038 | 0.2845 | 0.0842 | 0.7847 |
| MUNGED-FILTER | 0.2982 | – | – | 0.0955 | 0.0158 | 0.1190 | 0.5284 |
| CALL-GQ | 0.0463 | – | – | 0.0219 | 0.1071 | 0.1170 | 0.2923 |
| CALL-SB | 0.2840 | – | – | 0.0000 | 0.0000 | 0.0000 | 0.2840 |
| CALL-AF1 | 0.0299 | – | – | 0.0337 | 0.0232 | 0.1407 | 0.2275 |
| CALL-AD1 | 0.1194 | – | – | 0.0098 | 0.0208 | 0.0263 | 0.1763 |
| CALL-AF0 | 0.0040 | – | – | 0.0104 | 0.0434 | 0.0555 | 0.1134 |
| MUNGED-ID | 0.0377 | – | – | 0.0208 | 0.0068 | 0.0118 | 0.0771 |
| CALL-AD0 | 0.0025 | – | – | 0.0077 | 0.0347 | 0.0272 | 0.0722 |
| CALL-DPI | 0.0000 | – | – | 0.0059 | 0.0062 | 0.0340 | 0.0460 |
| INFO-MQ | 0.0014 | – | – | 0.0043 | 0.0032 | 0.0140 | 0.0230 |
| MUNGED-NEARBY | 0.0029 | – | – | 0.0023 | 0.0043 | 0.0083 | 0.0178 |
| INFO-SNVHPOL | 0.0037 | – | – | 0.0000 | 0.0000 | 0.0000 | 0.0037 |
| CALL-DP | 0.0012 | – | – | 0.0000 | 0.0000 | 0.0000 | 0.0012 |
| CALL-DPF | 0.0004 | – | – | 0.0000 | 0.0000 | 0.0000 | 0.0004 |

Table 12: This table shows the feature importances results for aligner "sentieon-201808.07" and caller "strelka-2.9.10". Importances are broken down by category with a cumulative sum at the end. Note that some results may be missing if the pipeline was run incorrectly or the models are not interpretable through `eli5`.

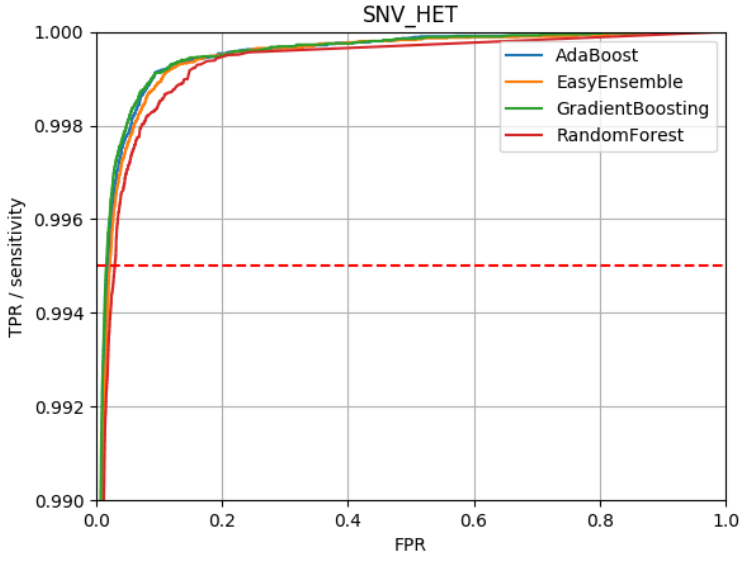

Figure 7: ROC curve for aligner "sentieon-201808.07", caller "strelka-2.9.10", variant type "SNV", and genotype "HET". Note that these curves are zoomed in to focus on only the region greater than the minimum clinical sensitivity (0.99).

###### 5.2.5 Model for SNV-HOM

Figure 8 contains the receiver-operator curves (ROC) for the final trained models for aligner "sentieon-201808.07", caller "strelka-2.9.10", variant type "SNV", and genotype "HOM".

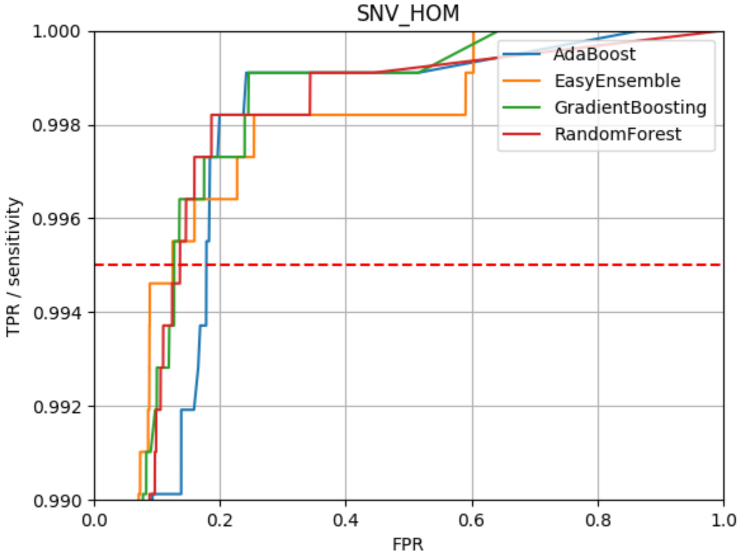

Figure 8: ROC curve for aligner "sentieon-201808.07", caller "strelka-2.9.10", variant type "SNV", and genotype "HOM". Note that these curves are zoomed in to focus on only the region greater than the minimum clinical sensitivity (0.99).

###### 5.2.6 Model for SNV-HE2

Figure 9 contains the receiver-operator curves (ROC) for the final trained models for aligner "sentieon-201808.07", caller "strelka-2.9.10", variant type "SNV", and genotype "HE2".

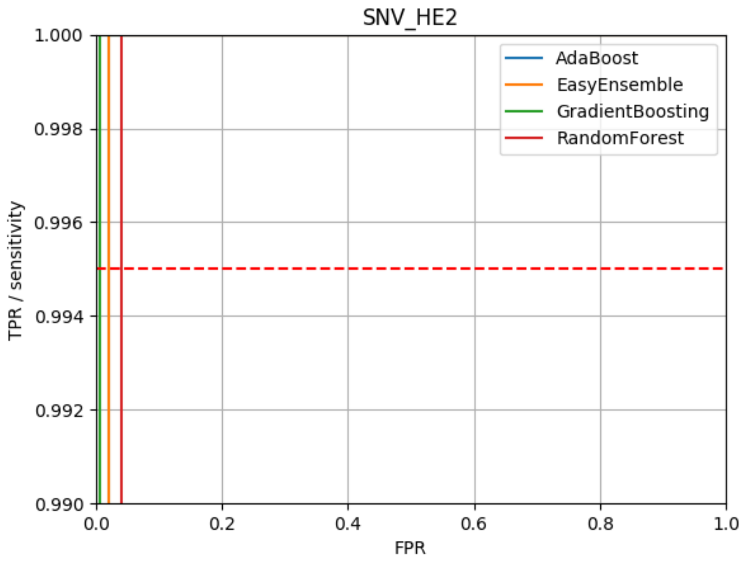

Figure 9: ROC curve for aligner "sentieon-201808.07", caller "strelka-2.9.10", variant type "SNV", and genotype "HE2". Note that these curves are zoomed in to focus on only the region greater than the minimum clinical sensitivity (0.99).

##### 5.2.7 Model for INDEL-HET

Figure 10 contains the receiver-operator curves (ROC) for the final trained models for aligner "sentieon-201808.07", caller "strelka-2.9.10", variant type "INDEL", and genotype "HET".

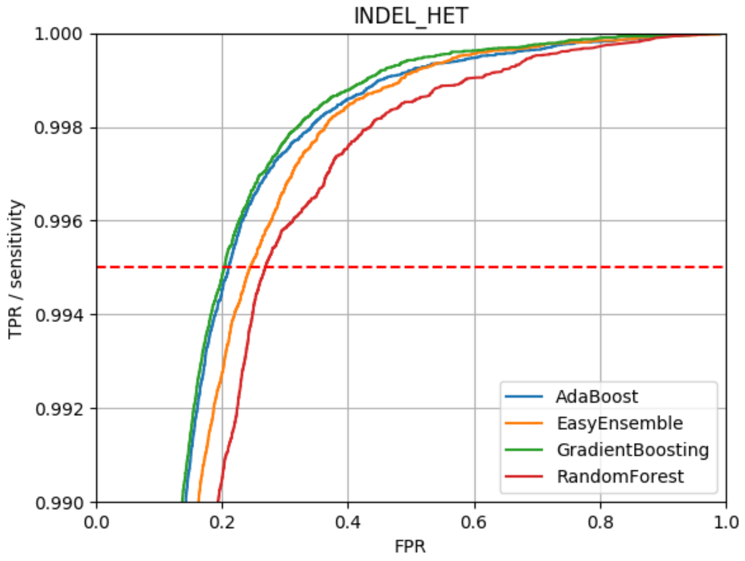

Figure 10: ROC curve for aligner "sentieon-201808.07", caller "strelka-2.9.10", variant type "INDEL", and genotype "HET". Note that these curves are zoomed in to focus on only the region greater than the minimum clinical sensitivity (0.99).

###### 5.2.8 Model for INDEL-HOM

Figure 11 contains the receiver-operator curves (ROC) for the final trained models for aligner "sentieon-201808.07", caller "strelka-2.9.10", variant type "INDEL", and genotype "HOM".

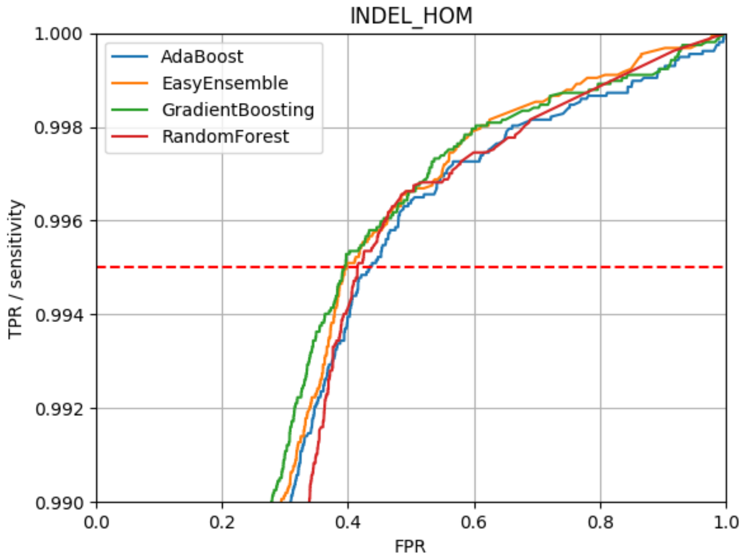

Figure 11: ROC curve for aligner "sentieon-201808.07", caller "strelka-2.9.10", variant type "INDEL", and genotype "HOM". Note that these curves are zoomed in to focus on only the region greater than the minimum clinical sensitivity (0.99).

###### 5.2.9 Model for INDEL-HE2

Figure 12 contains the receiver-operator curves (ROC) for the final trained models for aligner "sentieon-201808.07", caller "strelka-2.9.10", variant type "INDEL", and genotype "HE2".

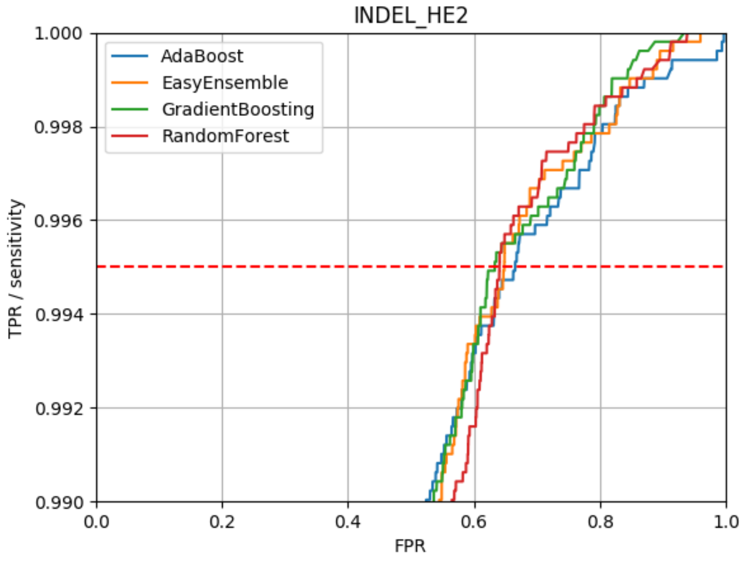

Figure 12: ROC curve for aligner “sentieon-201808.07”, caller “strelka-2.9.10”, variant type “INDEL”, and genotype “HE2”. Note that these curves are zoomed in to focus on only the region greater than the minimum clinical sensitivity (0.99).
